# Self-organizing Single-Rosette Brain Organoids from Human Pluripotent Stem Cells

**DOI:** 10.1101/2022.02.28.482350

**Authors:** Andrew M. Tidball, Wei Niu, Qianyi Ma, Taylor N. Takla, J. Clayton Walker, Joshua L. Margolis, Sandra P. Mojica-Perez, Roksolana Sudyk, Shannon J. Moore, Ravi Chopra, Vikram G. Shakkottai, Geoffrey G. Murphy, Jun Z. Li, Jack M. Parent

**Author notes:** Equal first authors (Andrew M. Tidball, Wei Niu). Corresponding Author: Jack M. Parent, MD, Address: Department of Neurology, 5021 BSRB, 109 Zina Pitcher Pl, Ann Arbor, MI 48109.

## Abstract

The field of brain organoid research is complicated by morphological variability with multiple neural rosette structures per organoid. We have developed a new human brain organoid technique that generates self-organizing, single-rosette spheroids (SOSRS) with reproducible size, cortical-like lamination, and cell diversity. Rather than patterning a 3-dimensional embryoid body, we initiate brain organoid formation from a 2-dimensional monolayer of human pluripotent stem cells (hPSCs) that is patterned with small molecules into neuroepithelium and differentiated to cells of the developing dorsal cerebral cortex. This approach recapitulates the 2D to 3D transition from neural plate to neural tube that occurs during neurodevelopment. The vast majority of monolayer fragments form spheres with a single central lumen and consistent growth rates. Over time, the SOSRS develop appropriately ordered lamination consistent with six cortical layers by immunocytochemistry and single cell RNA-sequencing. The reproducibility of this method has allowed us to demonstrate robust structural phenotypes arising from chemical teratogen exposure or when modeling a genetic neurodevelopmental epileptic disorder. This platform should advance studies of human cortical development, brain disorder mechanisms, and precision therapies.

**SUMMARY STATEMENT:** Simple procedure for generating reproducible single rosette cortical brain organoids used to identify robust structural phenotypes with neuroteratogen exposure and in a genetic neurodevelopmental disease model.

## INTRODUCTION

In 2008, Yoshiki Sasai first developed 3-dimensional cortical structures from mouse embryonic stem cells (ESCs) and later from human ESCs (hESCs) (Eiraku et al., 2008; Kadoshima et al., 2013). This method uses WNT and TGFβ inhibition to pattern an embryoid body (EB) into neuroepithelial tissue. Cells self-organize within the EB to form cortical neural rosette structures. These organizing centers are radially organized around a central lumen and represent *in vitro* correlates of neural tube formation during embryonic development. While embryos only have a single neural tube, these 3-dimensional (3D) *in vitro* structures contain many organizing centers. Other groups developed similar techniques to generate what began to be termed human *brain* or *cerebral organoids*, but in each case multiple rosettes form (Lancaster et al., 2013; Paşca et al., 2015). The variability of rosette formation is one of the key causes of a structural heterogeneity challenge in the brain organoid field (Quadrato and Arlotta, 2017). For this reason, many groups have used single-cell RNA-sequencing (scRNA-seq) to investigate development and disease as a non-structural read-out (Camp et al., 2015; Quadrato et al., 2017), while a majority of structural disease phenotypes observed to date in brain organoids reflect a size difference such as microcephaly or macrocephaly (Dang et al., 2021; Iefremova et al., 2017; Lancaster et al., 2013; Li et al., 2017; Qian et al., 2016). Although more detailed structural analyses are possible, the variability in size and shape of rosette structures necessitates both a high number of samples and a high level of observer selection (e.g. lumens on the outside edge of the organoid).

The structural heterogeneity and multiple organizing centers (neural rosettes) in brain organoids do not occur in other organoid systems. For example, intestinal (Spence et al., 2011) and pancreatic (Huang et al., 2015) organoids have a single lumen structure around which they organize. One main difference between these methods is that non-CNS organoids typically begin with patterning a 2-dimensional (2D) culture of pluripotent stem cells followed by 3D induction on a thick sheet of extracellular matrix (ECM) proteins. Similarly, *in vivo* neurodevelopment begins with patterning of the neural plate, a 2D structure, followed by neurulation to form the 3D neural tube. Therefore, we hypothesized that initiating organoid formation from a 2D cortical neuroepithelium (Chambers et al., 2009) would result in brain organoids with a single neural rosette organizing center.

We developed a protocol for cutting small, equal-sized fragments of dual-SMAD differentiated neuroepithelium and transferring them onto Geltrex, an ECM-like material. We found rapid self-organization of spheroids with single luminal centers expressing apical markers. We call these structures Self-Organizing Single Rosette Spheroids (SOSRS). Over time in culture, SOSRS recapitulate the normal inside-out lamination that occurs during cortical development and express markers of the normal complement of deep and superficial cortical layer neurons, followed by astrocytes. We show that the consistent size, architecture, and cell diversity of SOSRS at early timepoints allows for refined measurements for modeling both neural tube defect (NTD)-like chemical teratogenic effects and a genetic neurodevelopmental disorder caused by mosaic expression of protocadherin-19 (*PCDH19*).

## RESULTS

### 2D to 3D transition results in single rosette brain organoids

Our previous 2D neuronal differentiations of induced pluripotent stem cells (iPSCs) used 4 inhibitors to generate excitatory cortical-like neurons (Tidball et al., 2020). While most monolayer dual-SMAD inhibition protocols last 8-12 days (Chambers et al., 2009), we found consistent “rolling-up” and detachment of the monolayer between days 5-7 of differentiation. Therefore, we chose day 4 to initially test converting the 2D cultures to 3D by cutting the monolayer to reproducibly generate ∼125 µM squares that we replated onto a thick 100% Geltrex layer (**Fig. 1A-E**). Similar techniques have been used to induce 3D cultures since the 1980’s (Barcellos-Hoff et al., 1989). After 24 hours, the neuroepithelial squares on ECM rounded up into spheres with a central apical lumen (**Fig. 1D** and **Movie S1**). Live-imaging of SOSRS formation with the apical lumen labelled with zonula occludens-1 (ZO-1)-EGFP demonstrated fluorescence migration from the edges to a central lumen in 6 hours, a finding consistent with a neurulation-like event (**Movie S2**). The efficiency of single lumen formation across 3 control human iPSC lines was 84 ± 9 % at 6 days *in vitro* (**Fig. 1F,G,I**). However, SOSRS with 2 lumens had an overall area that was 2.2 times the average for single lumen SOSRS (**Fig. 1H**) and a linear regression of the relationship between lumen number and SOSRS area has a slope of 13,500 µm^2^ while the average area of single lumen SOSRS was 12,400 µm^2^ (**Fig. 1J**). These data suggest that multiple lumens are caused by the fusion of multiple monolayer fragments and that the single neural rosette/lumen efficiency for individual fragments is higher than 84%.

**Figure 1.**
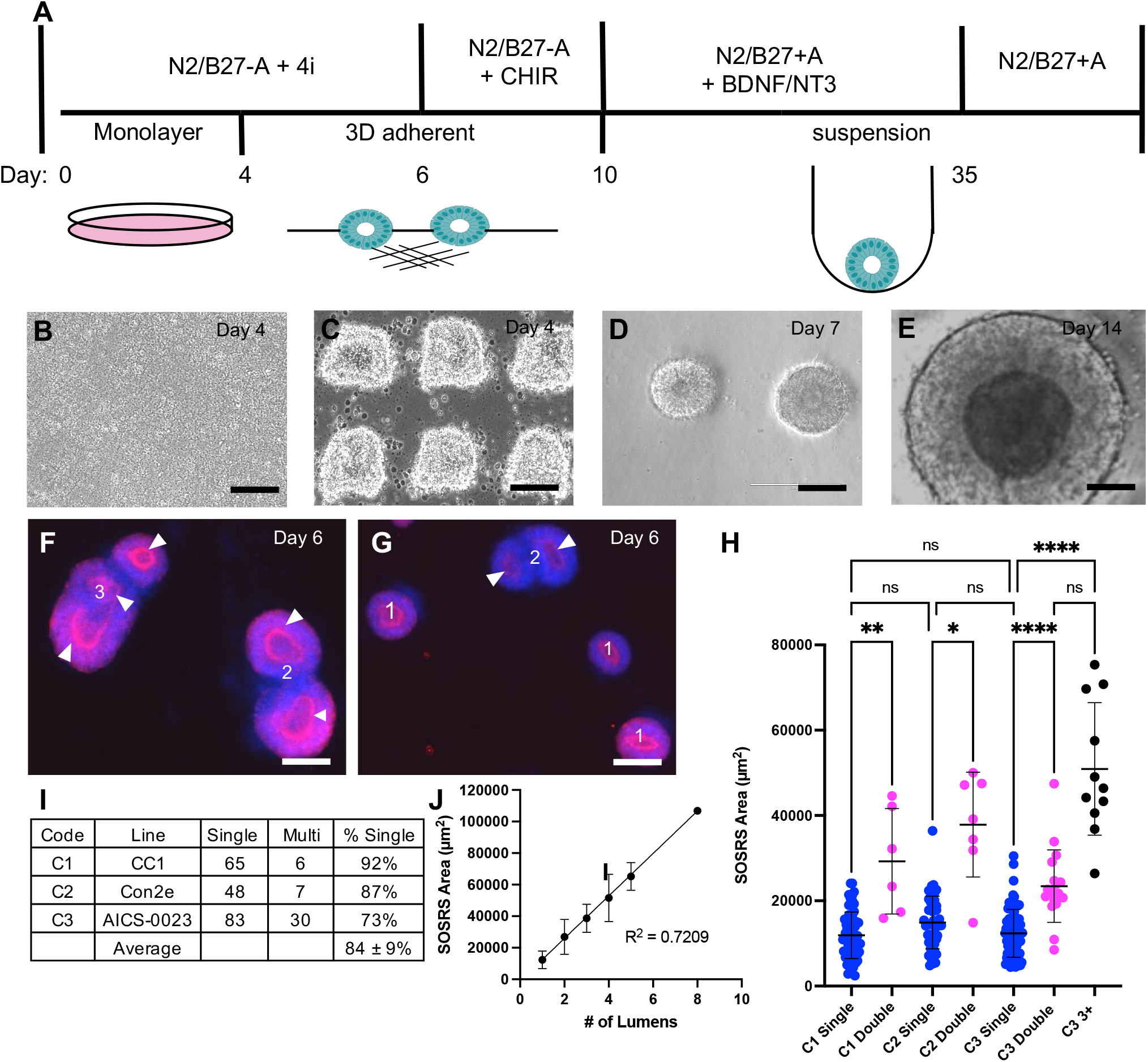
Two-dimensional to three-dimensional transition results in majority single rosette spheroids. (**A**) Schematic of SOSRS differentiation timeline. The top half describes the media components while the bottom half is the culture format. **B-E** Phase micrographs of SOSRS at important stages including neuroepithelial monolayer (**B**), monolayer cutting (**C**), early SOSRS formation on extracellular matrix (**D**), and a SOSRS in suspension at day 14 (**E**). (**F**,**G**) Representative images of ZO-1 immunostained SOSRS with bis-benzamide counter stain depicting single (1), double (2), and triple (3) lumens in SOSRS with obvious fusion of multi-rosette SOSRS. For multi-rosette SOSRS, each lumen is identified by an arrowhead. (**H**) Areas of SOSRS with one (single or 1), two (double or 2) or more (3+) lumens for each iPSC line (C1, C2, and C3). * = p < 0.05, ** = p < 0.01, and **** = p < 0.0001. (**I**) Table of SOSRS numbers with single or multiple lumens and the percentages with single lumens from three different iPSC lines, as well as the mean overall percentage of SOSRS with single lumens. (**J**) XY plot of all lumen data with lumen # on the X-axis and SOSRS size on the Y-axis. Linear regression was performed, and the goodness of fit R^2^ = 0.7209. Error bars are s.d. All scale bars are 100 µm.

We next assessed SOSRS for the development of early ventricular zone (VZ), subventricular zone (SVZ) and cortical mantle-like structures. Day 8 SOSRS show homogeneous neuroepithelial marker expression of PAX6 and nestin with radially organized tubulin networks (**Fig. 2A,B**). The inner lumens label with the apical marker PKC-z (**Fig. 2B**) demonstrating correct polarization, and neural stem cells displayed potential interkinetic nuclear migration with mitotic cells positioned on the apical surface (**Fig. 2B, arrowhead**). On day 7, the SOSRS showed significant peripheral cell death, but the addition of CHIR99021 from days 6-10 blocked much of the cell death (cleaved caspase-3 staining) and allowed rapid SOSRS growth (**Fig. S1A**,**B**). CHIR99021 exposure yielded SOSRS of significantly larger SOSRS size and decreased intensity of DRAQ7, a non-cell permeable DNA dye used to measure cell death (**Fig. S1C**,**D**). A similar CHIR99021 pulse has been used previously in human brain organoid cultures to expand rosette size (Lancaster et al., 2017), and likewise, this pulse in SOSRS resulted in rapid expansion of the PAX6-positive cells while maintaining a large central lumen (**Fig. 2C,D**).

**Figure 2.**
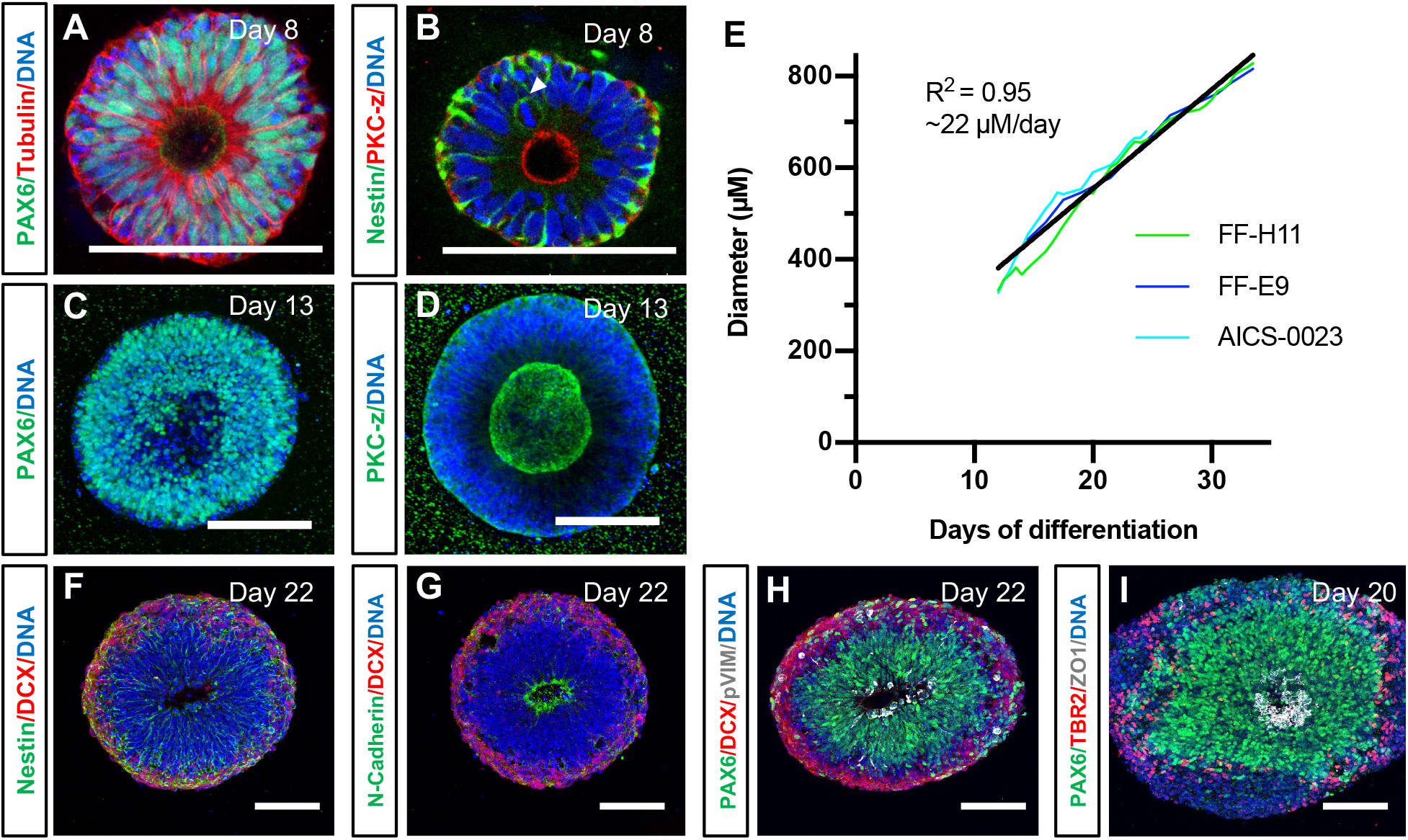
SOSRS demonstrate characteristics of early cortical development and consistent growth kinetics. (**A-D**) Confocal micrographs of whole mount day 8 (**A**,**B**) or day 13 (**C**,**D**) SOSRS immunostained for the designated proteins. (**E**) SOSRS generated from 3 different iPSC lines were grown in the Incucyte live-cell imaging system with average diameters plotted and used to generate the growth curve slope and R^2^. FF-H11: n = 11; FF-E9: n = 10; AICS-0023: n = 60. (**F-I**) Confocal micrographs of cryosectioned SOSRS immunostained for neural markers on days 20-22. Nuclei were stained with bis-benzamide (DNA, blue) for all immunofluorescence images, and days of differentiation are indicated in the upper right corner of each image. All scale bars are 100 µm.

### Continued SOSRS maturation after removal from ECM

To allow continued expansion, we removed the SOSRS from Geltrex between days 9-11. A diameter of ∼ 250 µM was optimal for cell survival and maintenance of the single rosette at this stage (data not shown). The SOSRS were transferred to low-attachment 96-well plates for suspension culture with CHIR99021 included for the first 24 hours to increase cell survival. Brain-derived neurotrophic factor (BDNF) and neurotrophin-3 (NT3) were added to the culture medium from this time until day 35. The 96-well plates allowed for the isolation of individual SOSRS to avoid unintentional fusion. Daily SOSRS live imaging enabled longitudinal growth monitoring and demonstrated growth kinetic reproducibility between lines and batches (**Fig. 2E**).

### SOSRS demonstrate lamination characteristic of human cortical development

Day 22 SOSRS displayed an evenly distributed layer of neurogenesis at the periphery as shown by the expression of mCherry driven by the doublecortin (DCX) promoter (**Fig. 2F-H**). A VZ-like region persisted in the interior adjacent to the lumen that consisted of PAX6-positive cells with phospho-vimentin labeled dividing radial glia (RG) on the lumen surface (**Fig. 2H**). A second proliferative zone appeared on the outside of the organoid and likely reflected dividing intermediate progenitors (IPs) (**Fig. 2H**) as the IP marker TBR2 (*EOMES*) labeled a ring of cells in this same relative position (**Fig. 2I**). The SOSRS continued to show a clear layer of TBR2 expressing cells just outside the PAX6-expressing VZ at six weeks in culture, similar to the TBR2+ SVZ of the developing cortex (**Fig. 3A-C**), and these cells were detected by scRNA-seq up to at least 5 months (**Fig. 6R**).

**Figure 3.**
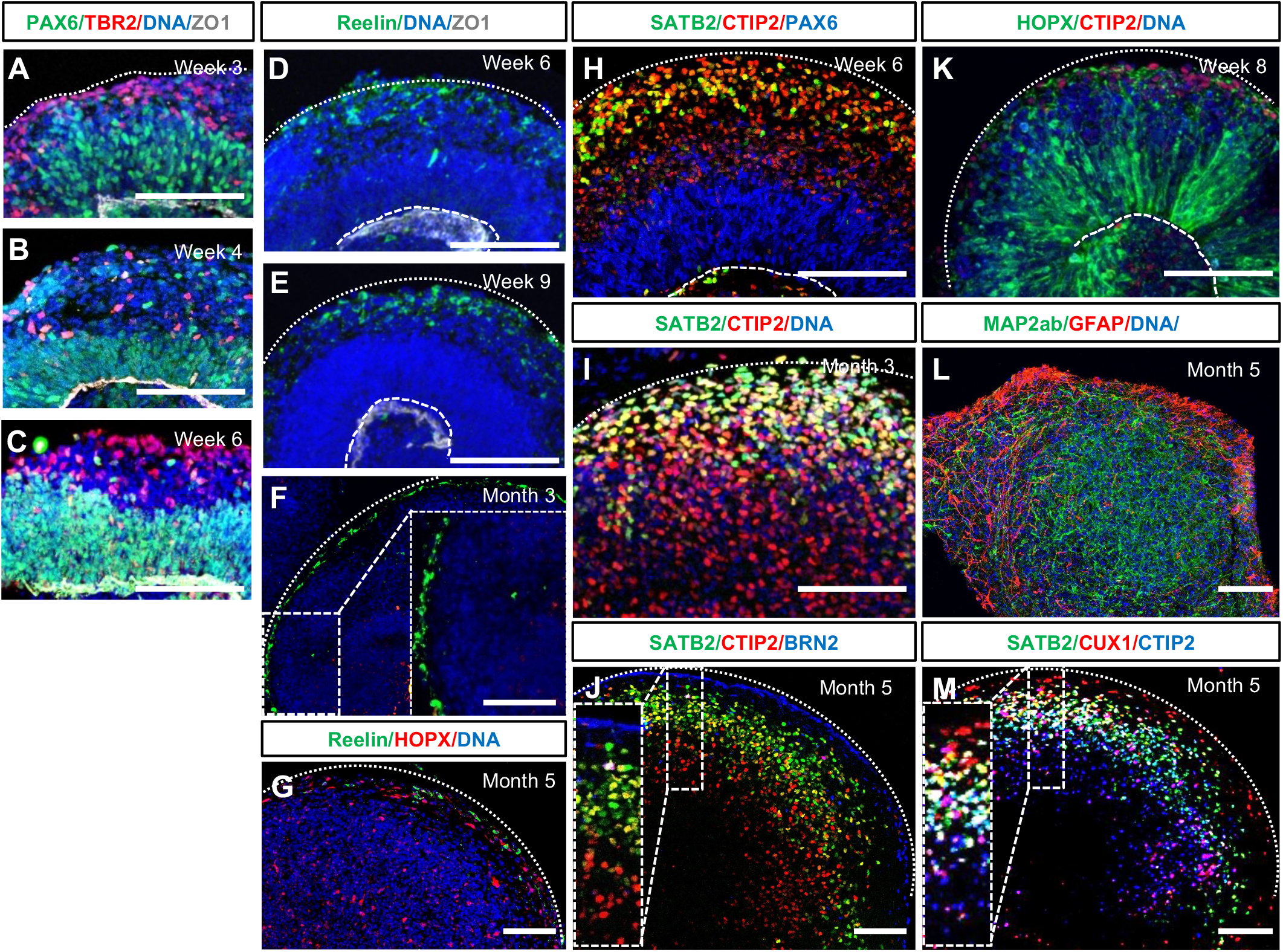
SOSRS have neurodevelopmentally consistent cortical lamination patterns. (**A-C**) SOSRS immunostained for the radial glial marker PAX6 (green) and intermediate progenitor marker TBR2 (red). ZO-1 immunolabeling (white) outlines the apical lumen, and bis-benzamide nuclear stain (DNA) is in blue. (**D-G**) SOSRS at several timepoints immunostained for Reelin (green) and HOPX (red; **G** only). Bis-benzamide nuclear stain (DNA) is in blue. (**H-J**,**M**) SOSRS immunostained for cortical layer markers CTIP2, SATB2, BRN2, or CUX1. The boxed areas in (**F**,**J**,**M**) are magnified in the insets. (**K**) 8-week SOSRS begin to express the oRG marker HOPX with radially oriented processes. (**L**) SOSRS at 5 months express the astrocyte marker GFAP on the outer edge and the mature neuronal marker MAP2ab throughout. Dashed curved lines in (**D-E**,**H**,**K**) mark the apical surface of the lumen, and dotted curved lines present in most panels denote the outer edge of the SOSRS. Scale bars are 100 µm.

Neuronal layering of the SOSRS followed the typical pattern for cortical development. First, Reelin+ cells were found in the most superficial layer by week 6, persisted for at least 5 months, and remained located on the SOSRS periphery (**Fig. 3D-G**), analogous to Reelin+ Cajal-Retzius cells located in the marginal zone of the developing cortex. Reelin+ cells also appeared in one-, three-, and five-month scRNA-seq data (**Figs 4X, 5B’, 6S**). By 6 weeks, we observed many cells expressing the deep layer cortical marker CTIP2 (*BCL11B*) peripheral to the PAX6+ proliferative zone with more superficial layer SATB2+ cells intermingled (**Fig. 3H**). In three-month SOSRS, more SATB2+ cells appeared superficial to the CTIP2 layer but with continued areas of overlap (**Fig. 3I**). After 5 months, SOSRS show clear separation of CTIP2+ and SATB2+ cell layers, similar to other organoid protocols and developing human fetal brain (Qian et al., 2020), with robust expression of the upper layer neuronal markers BRN2 and CUX1 peripheral to the SATB2+ layer (**Fig. 3J,M**). Therefore, our model shows developmentally regulated timing and positioning of cortical layer markers consistent with *in vivo* human cortical development. Notably, we only observed neurodevelopmentally normative layering in SOSRS displaying continued expression of Reelin+ cells on their exterior, consistent with the importance of these cells in normal “inside-out” cortical development (Ogawa et al., 1995).

**Figure 4.**
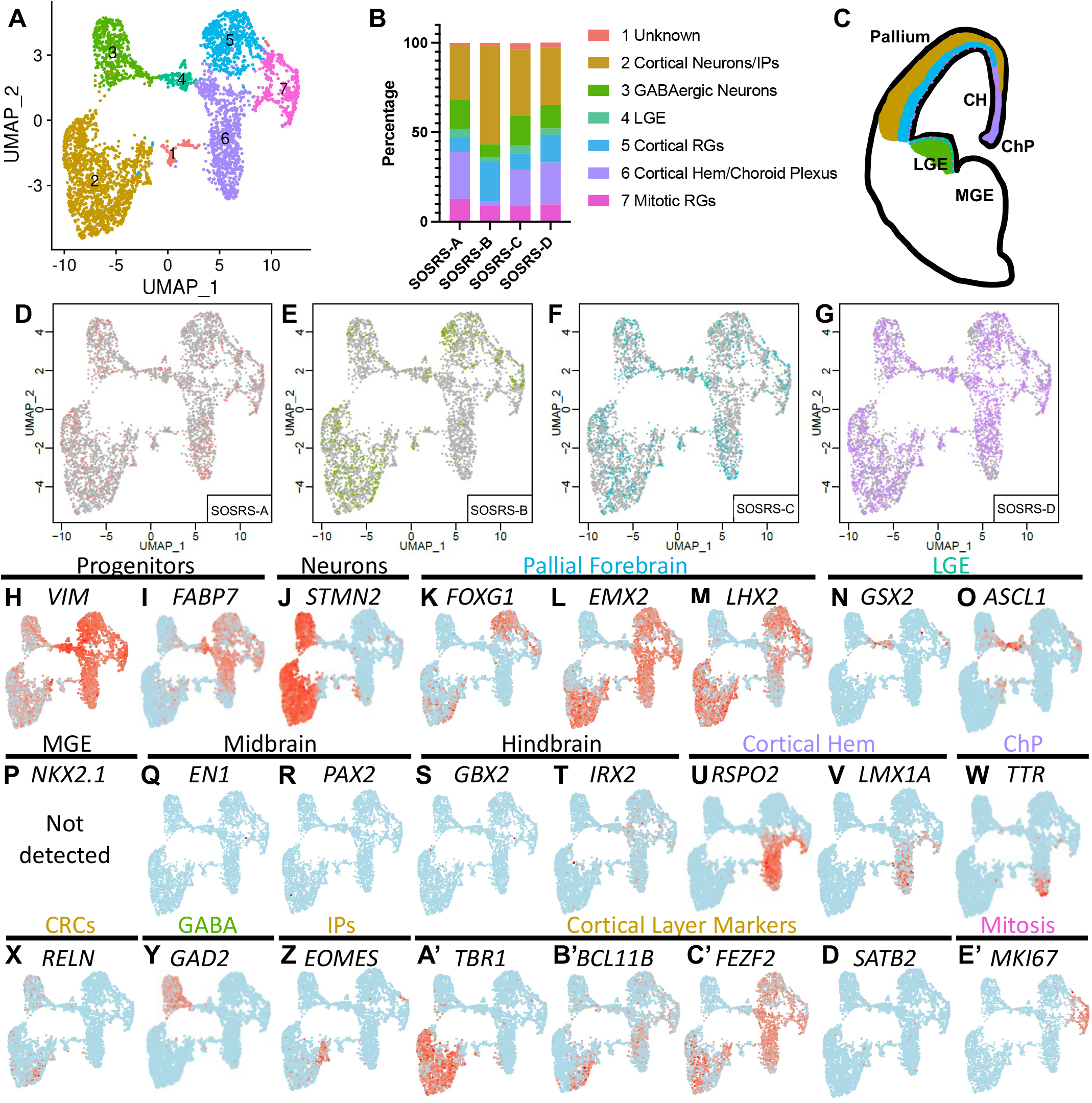
Single cell RNA sequencing for 4 separate one-month SOSRS from line AICS-0023. (**A**) UMAP plots for one-month SOSRS with color coding of identified clusters. (**B**) Bar graph showing the percentage of cells found in each cluster of the 4 one-month SOSRS. The legend at right displays the cell type identity for each cluster. (**C**) Cartoon of developing telencephalon with the color-coded identity of each cluster labeled. (**D-G**) UMAP plots for each of the four one-month SOSRS. Colored dots are for the depicted SOSRS while gray dots are from the other 3 SOSRS. (**H-E’**) Individual UMAP plots of selected genes from each of the 4 one-month SOSRS. The region each marker identifies is listed above the gene names. Regions that are cluster identities are color-coded to match panels (**A-C**).

**Figure 5.**
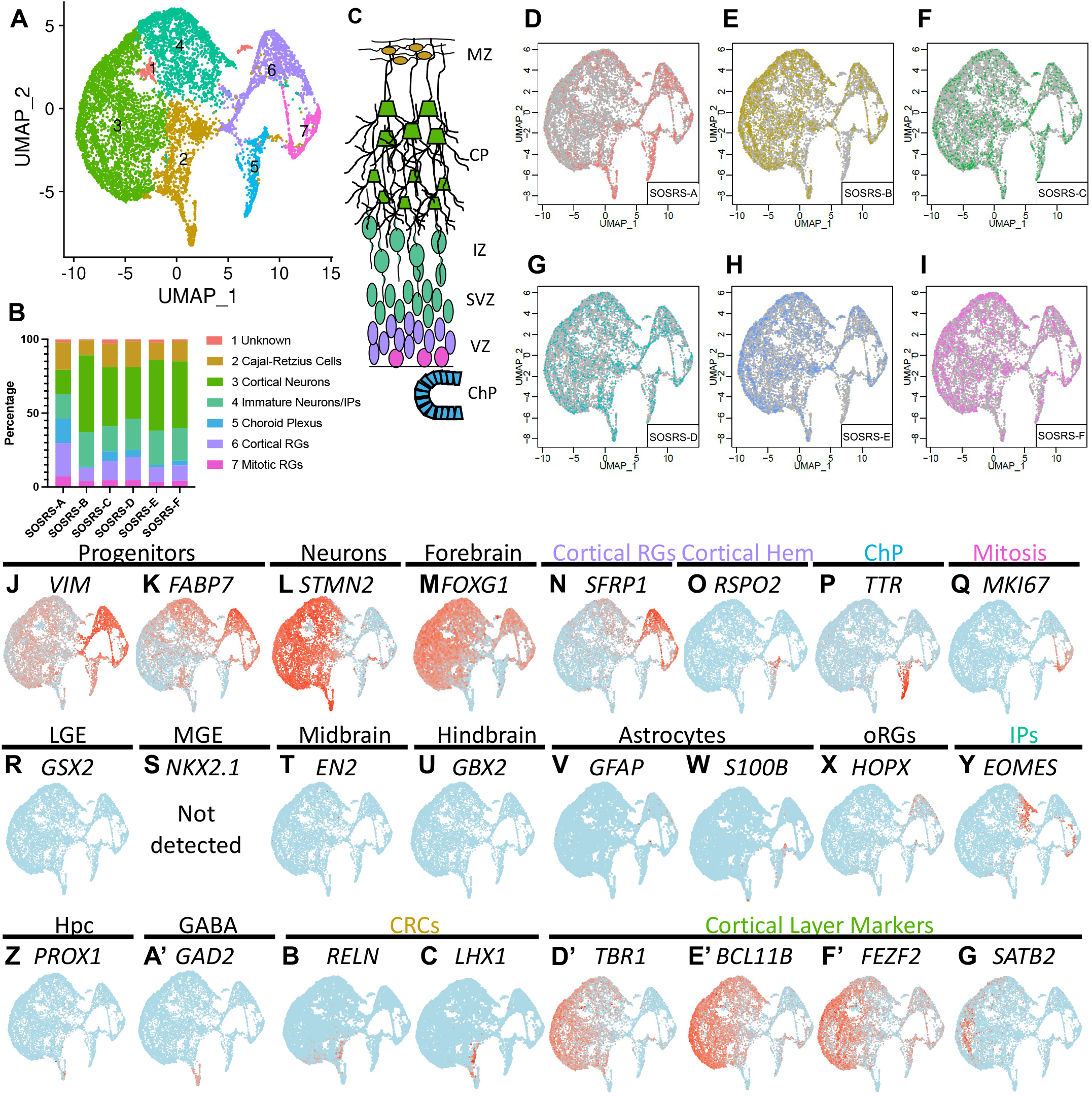
Single cell RNA sequencing for 6 separate three-month SOSRS from line AICS-0023. (**A**) UMAP plots for three-month SOSRS with color coding of identified clusters. (**B**) Bar graph showing the percentage of cells found in each cluster of the 6 three-month SOSRS. The legend at right displays the cell type identity for each cluster. (**C**) Cartoon of developing dorsal telencephalon layers with the color-coded identity labeled for each cluster found in three-month SOSRS. (**D-I**) UMAP plots for each of the 6 three-month SOSRS. Colored dots are for the depicted SOSRS while gray dots are from the other 5 SOSRS. (**J-G’**) Individual UMAP plots of the specified genes for each of the 6 three-month SOSRS. The region each marker identifies is listed above the gene names. Regions that are cluster identities are color-coded to match panels (**A-C**).

**Figure 6.**
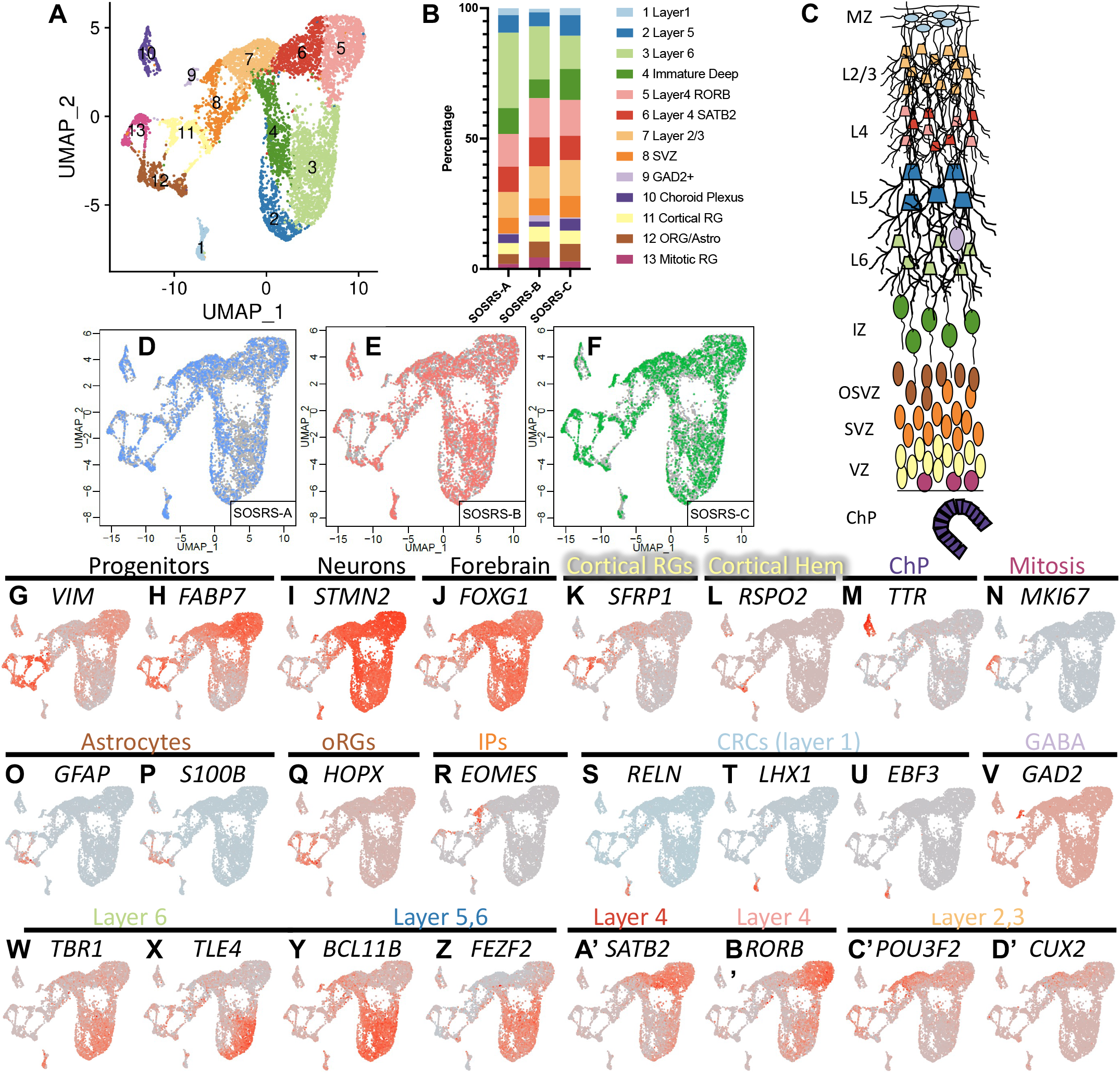
Single cell RNA sequencing for 3 separate five-month SOSRS from line AICS-0023. (**A**) UMAP plots for five-month SOSRS with color coding of identified clusters. (**B**) Bar graph showing the percentage of cells found in each cluster of the 3 five-month SOSRS. The legend at right displays the cell type identity for each cluster. (**C**) Cartoon of developing dorsal cortex layers with the color-coded identity of each cluster labeled. (**D-F**) UMAP plots for each of the 3 five-month SOSRS. Colored dots are for the depicted SOSRS while gray dots are from the other 2 SOSRS. (**G-D’**) Individual UMAP plots of the specified genes for each of the 3 five-month SOSRS. The region each marker identifies is listed above the gene names. Regions that are cluster identities are color-coded to match panels (**A-C**).

The presence of superficial cortical neurons should follow the appearance of outer radial glia (oRG) which primarily give rise to these neuronal subtypes in primate development (Nowakowski et al., 2016). Indeed, HOPX+ cells, a quintessential oRG marker, arose in two-month SOSRS (**Fig. 3K**). HOPX expression was minimal in one-month SOSRS by scRNA-seq but was present in ∼30% of RG by 3 months and persisted for at least 5 months (**Figs 3G, 6Q**). Astrogenesis occurs very late in the developing cortex, and, likewise, we found little evidence of astrocytes present in three-month SOSRS by scRNA-seq (**Fig. 5V,W**). We did observe robust expression of the astrocyte marker GFAP in five-month SOSRS, mostly on the periphery (**Fig. 3L**). While a single rosette/lumen structure persisted during the first 9 weeks of SOSRS differentiation (**Figs 1, 2**), after this timepoint many SOSRS developed multiple rosettes or lost all rosette structure (data not shown); however, our cortical layer marker expression demonstrated that normal developmental layering still occurred. We next examined whether neurons in three- or five-month SOSRS were electrophysiologically active. First, dissociated neurons from 3-month SOSRS were plated onto glass coverslips, labeled with lentiviral calcium/calmodulin-dependent protein kinase II-alpha (CAMKIIα) and cultured for five weeks. Whole-cell patch-clamp recordings showed that the neurons had depolarized resting membrane potentials (−30 ± 5 mV) but were able to fire trains of action potentials when injected with pulses of depolarizing current (**Fig. S2**). We next cultured SOSRS on multielectrode array (MEA) recording plates at ∼ 100 days and recorded for approximately 8 weeks. The SOSRS showed an expected increase in action potential firing and burst activity over time (**Fig. S3**). Therefore, neurons from SOSRS are electrophysiologically active – albeit immature – at 3 months and show increasing activity through at least 5 months.

### SOSRS have reproducible cell type diversity

To further characterize cell types and variability between individual SOSRS, we performed scRNA-seq of SOSRS at one, three, and five months using 4, 6, and 3 SOSRS per time point, respectively. For all scRNA-seq experiments, the commercially available AICS-0023 iPSC line was used because of its robust validation. The number of post QC cells used for analysis for each SOSRS are listed in **Table S1**. For the 4 one-month SOSRS, we live-imaged from days 9-31 (**Movie S3**) and found that average growth kinetics were indistinguishable from 38 additional organoids imaged in the same batch (**Fig. S4A**). At each of the three timepoints, UMAP plots formed distinct clusters that were reproducible between individual SOSRS (**Figs 4-6**). One-month SOSRS formed 7 distinct gene expression clusters that were well reproduced in the 4 separate SOSRS except for minor differences in SOSRS-B (**Fig. 4A-G**). Similarly, three-month SOSRS separated into 7 distinct cell type clusters with remarkable consistency across the 6 different SOSRS, although some distinct differences in SOSRS-A arose with more cells in progenitor clusters and fewer in neuronal clusters (**Fig. 5A-I**). The five-month SOSRS had a greater number of cell type clusters, 13, with continued reproducibility between the 3 individual SOSRS (**Fig. 6A-F**). Cross-correlation analysis of each cluster between samples at the one- or three-month timepoints show a remarkable level of transcriptomic similarity when using the top 50 markers of each cluster (**Fig. S4B**,**C**), except in the unidentified cell cluster 1. Thus, both the population of each cell type and the transcriptomes within each cell type cluster are remarkably similar between individual SOSRS.

To identify the cell types in each cluster, we first relied on markers of RG progenitors (*VIM* and *FABP7*) and neurons (*STMN2*) (**Figs 4H-J, 5J-L, 6G-I**). Almost all RG (clusters 5-7, **Fig. 4A**) expressed the dorsal forebrain markers *EMX2* and *LHX2* (**Figs 4L-M)**, but only a portion (clusters 5 and 7) expressed *FOXG1* (**Fig. 4K**), a standard forebrain marker. We checked for ventral forebrain marker expression and found that Cluster 4 expressed markers of the lateral ganglionic eminence (LGE), *GSX2* and *ASCL1*, while there was no expression of the medial ganglionic eminence (MGE) marker, *NKX2*.*1*, in our dataset (**Fig. 4N-P**). We also observed almost no expression of midbrain (*EN1* and *PAX2*) or hindbrain markers (*GBX2* and *IRX2*) (**Figs 4Q-T, 5T-U**). The cortical hem, while a part of the dorsal forebrain, expresses little or no *FOXG1* or *PAX6*, but does express *EMX2* and *LHX2* (De Clercq et al., 2018; Shinozaki et al., 2004). We confirmed the hypothesis that the *FOXG1*-negative cluster 6 is cortical hem by observing clear expression of *RSPO2*, a cortical hem specific marker (La Manno et al., 2021), and the roof plate marker *LMX1A* (**Fig. 4U,V**). The cortical hem is a major contributor of Cajal-Retzius cells (Meyer, 2010); therefore, these progenitors are likely the source of the *RELN-*expressing cells found by immunostaining (**Fig. 3D-G**) and scRNA-seq (**Figs 4X, 5B’, 6S**).

The *RELN-*expressing clusters at three- and five-months expressed other Cajal-Retzius markers, including *LHX1* and *EBF3* (**Figs 5C’, 6T-U**). Cells expressing the choroid plexus marker transthyretin (*TTR*) were seen at one- and three-months in the same cluster as the cortical hem (**Figs 4W, 5P**). The choroid plexus anlage is found next to the cortical hem in the developing cortex; therefore, expression of other transcripts may be similar between these cell types leading to a shared cluster. In five-month SOSRS, the *TTR* expressing choroid plexus cells make their own distinct cluster. Cluster 7 cells in one-month SOSRS express markers of cortical RG, and cell cycle stage-specific transcript analysis determined that it is defined by markers of M and S phase cells (**Fig. S5**) as shown by the mitotic marker *MKI67* (**Fig. 4E’**). We color-coded the regional identity of each cluster on a map of the developing telencephalon in **Fig. 4C**.

Within the neuronal clusters, we expected mainly excitatory cortical neurons as shown by the marker *TBR1* (**Fig. 4M**). However, about 25% of the early-born neurons in one-month SOSRS were GABAergic neurons (as defined by *GAD2* and *DLX2* (**Fig. 4Y** and **data not shown**). These likely originate from the LGE-like dividing cells expressing *GSX2* (**Fig. 4N**). At three- and five-months, however, *GAD2*+ cells account for less than 1% of all cells (**Figs 5A’, 6V**). Therefore, while some LGE-derived neurons exist in our SOSRS, they are a very small fraction at later timepoints. TBR2/*EOMES* expressing intermediate progenitors (IPs) (**Fig. 4Z**) did not resolve into their own cluster at any of the timepoints but were grouped with cells expressing markers of immature, migrating neurons including *BHLHE2*2, *NHLH1, NeuroD1*, and *NeuroD6* (See **Tables S2-4**). Interestingly, IPs are located in the only region of their respective clusters that are negative for the pan-neuronal marker *STMN2* (**Figs 4J,Z, 5L,Y, 6I,R**).

Cortical layer specific markers appeared in distinct clusters following typical developmental timing. At one-month, layer 1 *RELN* expressing cells and deep layer *TBR1, BCL11B* (CTIP2), and *FEZF2* neurons were present (**Fig. 4X, A’-C’**). Layer 4 markers *SATB2* and *RORB* appeared at three- and five-months, respectively (**Figs 5G’, 6A’,B’**). The superficial, layer 2,3 neuronal markers *POU3F2* (BRN2) and *CUX2* are only present in five-month SOSRS (**Fig. 6C’,D’**). Astrocytes are the last cell type produced by dorsal RG in the developing cortex. Accordingly, markers of astrocytes, including *GFAP* and *S100B*, are not present until five-months (**Fig. 6O,P**). This coincides with the increase in *HOPX* expressing outer radial glial cells known to give rise to superficial neurons and astrocytes in the human cortex (**Fig. 6Q**). The progression over time is also visualized by dot matrix plots of selected genes (**Fig. S6**). Therefore, by five-months in culture, most cell types descendent from the RG of the pallial VZ and cortical hem are present in SOSRS and appear in a typical neurodevelopmental progression.

### SOSRS offer a platform for studying NTDs

We next explored the translational potential of SOSRS for studying human neurodevelopmental disorders. NTDs are a potential teratogenic adverse consequence of pharmaceuticals used during pregnancy (Ornoy, 2006). Proper neural tube formation depends upon the apical constriction of neuroepithelial cells (**Fig. 7A**), a process that occurs via the canonical SHROOM3/Rho-kinase/non-muscle myosin II pathway (Hildebrand, 2005). To determine if this mechanism underlies SOSRS formation during 2D to 3D conversion, we explored the effects of adding a Rho-kinase inhibitor, Y-27632, or a non-muscle myosin inhibitor, blebbistatin, to the culture medium on day 5 of differentiation. SOSRS were examined 24 hours later for the tight-junction marker ZO-1 and microtubule network marker acetylated-tubulin. In vehicle-treated SOSRS, the tight-junction network formed a spherical lumen with small apical surfaces and a radially oriented microtubule network (**Fig. 7B,F)**. Both inhibitor-treated SOSRS, in contrast, displayed disorganization of the tight-junctions and microtubules (**Fig. 7C,G**). Quantification of several SOSRS metrics showed increased apical endfeet surface area after Y-27632 treatment (**Fig. 7E**), and a larger lumen/total SOSRS surface area ratio induced by exposure to either inhibitor (**Fig. 7D,H**). Additionally, lumen circularity was reduced by blebbistatin (**Fig. 7I**). Incubation with Y-27632 on day 4 when the monolayer fragments were first placed on ECM completely blocked lumen formation (data not shown). These findings show that apical constriction is important for SOSRS formation and suggest that they follow the normal mechanism of neurulation.

**Figure 7.**
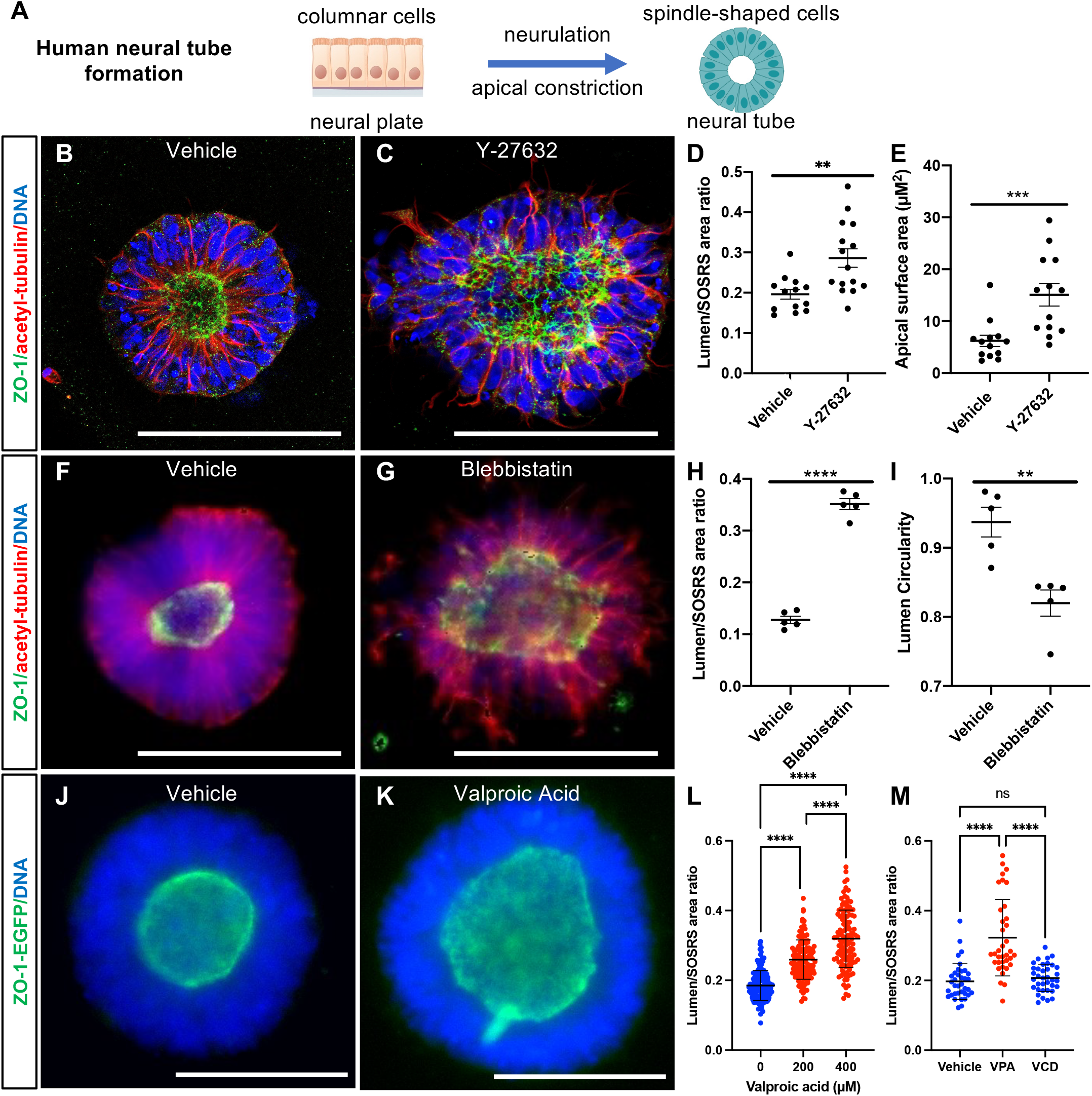
Teratogenic compound exposure results in dysmorphic SOSRS with enlarged lumens. (**A**) Schematic of cellular and structural changes during neurulation. (**B**,**C**,**F**,**G**,**J**,**K**) Confocal (**B**,**C**) or epifluorescence (**F**,**G**,**J**,**K**) micrographs of whole-mount SOSRS either immunostained for ZO-1 (green) and acetyl-tubulin (red) (**B**,**C**,**F**,**G**), or expressing ZO-1-EGFP fusion protein (**J**,**K**). All images show nuclear bis-benzamide stain (blue). SOSRS were exposed to vehicle (untreated; **B**,**F**,**J**) or inhibitors of the apical constriction pathway, either Y-27632 (**C**, 20 µM), blebbistatin (**G**), or the teratogenic antiseizure medication, valproic acid (**K**, 200 µM). (**D**,**E**,**H**,**I**,**L**,**M**) Quantification of structural measurements including apical surface area of individual endfeet (**E**), lumen area/SOSRS area (**D**,**H**,**L**,**M**) or lumen circularity (**I**). (**L**) Data points are individual SOSRS from 6 independent experiments with 0, 200, and 400 µM valproic acid. Each individual experiment also showed a significant increase for both 200 and 400 µM valproic acid. n = 139, 127, and 123, respectively. (**M**) Data points are individual SOSRS from 3 independent experiments, n = 36 for each group. Welch’s t-test was used for two-group comparisons, and Kruskal-Wallis test was used for more than two groups. ** = p < 0.01, *** = p < 0.001, and **** = p < 0.0001. Error bars are s.e.m. for (**D**,**E**,**H**,**I**) and s.d. for (**L**,**M**). Scale bars are 100 µm.

After establishing this methodology, we applied the SOSRS platform to a well-established neuroteratogenic drug, valproic acid. Valproic acid has been used for decades as an effective antiseizure medication, but it is also a potent neuroteratogen with a high risk of NTDs arising from pregnancies in women treated with valproic acid (Robert and Guibaud, 1982). ZO-1-EGFP fusion protein-expressing SOSRS exposed to valproic acid from days 4-6 showed a dose-dependent increase in normalized lumen area at 200 µM and 400 µM doses in each of 6 independent experiments and as a combined data set (**Fig. 7L**). Representative images near the mean values for vehicle and 200 µM valproic acid treatment are shown in **Fig. 7J,K**. Lastly, in 3 additional independent experiments we treated with either 400 µM valproic acid or 400 µM valnoctamide, a valproic acid derivative with similar antiseizure activity but no teratogenicity in mice (Radatz et al., 1998). In these experiments, valproic acid significantly increased the lumen area but valnoctamide had no effect, indicating that the SOSRS are modeling the teratogenic mechanism of valproic acid. Therefore, these findings suggest that SOSRS are a useful platform for modeling human NTDs.

### SOSRS recapitulate abnormal cell segregation associated with mosaic *PCDH19* expression

We next explored whether SOSRS are useful for modeling alterations of the developing cortex in a genetic neurodevelopmental disorder. Protocadherin-19 (PCDH19) clustering epilepsy (PCE) is a severe childhood developmental and epileptic encephalopathy caused by inherited or *de novo*, heterozygous loss-of-function variants in the X-linked *PCDH19* gene (Dibbens et al., 2008; Kolc et al., 2019). Intriguingly, pathogenic variants only lead to epilepsy and intellectual disability in heterozygous females and mosaic males, but not in hemizygous mutant males who are asymptomatic carriers (de Lange et al., 2017). A prevailing hypothesis for this phenomenon is that disease manifestations reflect the mosaic pattern of wild-type (WT) and mutant *PCDH19*-expressing cell populations in the developing brains of heterozygous females caused by random X-inactivation, or in mosaic males. Because PCDH19 is involved in homophilic cell-cell interactions during development (Hoshina et al., 2021; Pederick et al., 2018), the inability of WT and mutant *PCDH19*-expressing cells to interact is thought to cause disease. This hypothesis is supported by findings in female heterozygous *PCDH19* knockout (KO) mice in which mutant and WT *PCDH19*-expressing cells segregated into distinct columns in the VZ, SVZ, and cortex (Hoshina et al., 2021; Pederick et al., 2018).

To determine whether mosaic cell segregation can be modeled *in vitro* using human brain organoids, we first generated male lines with frameshift mutations in *PCDH19*, along with isogenic controls, by simultaneous CRISPR gene editing and iPSC reprogramming (Tidball et al., 2017; Tidball et al., 2018). We confirmed loss of *PCDH19* mRNA expression in the KO lines by qRT-PCR (**Fig. S7A**,**B**). Each KO and isogenic control line was then GFP-labeled by stable integration. To model a PCE mosaic male, equal amounts of single-cell dissociated EGFP-labeled WT and unlabeled *PCDH19* KO iPSCs were mixed one day prior to beginning organoid differentiation. For comparison, mixtures of labeled and unlabeled cells of the same genotype (WT:WT or KO:KO) were used to generate organoids (**Fig. 8A-C**). Both SOSRS and multi-rosette brain organoids using the spin-Ω method (Qian et al., 2016) were generated in parallel. At day 20, mixed WT/*PCDH19* KO SOSRS (**Fig. 8G**), but not spin-Ω cortical organoids (**Fig. 8D**), showed robust cell segregation that was not evident in any of the non-mosaic WT/WT or KO/KO mixed conditions using either brain organoid method (**Fig. 8E,F**,**H**,**I**). This robust phenotype was consistent across multiple SOSRS (**Fig. 8J,K**), and was reproduced in multiple batches (**Fig. S7C**,**D**) and with a female hESC line (**Fig. S7E-G**). These data indicate that a structural developmental brain abnormality associated with a genetic neurodevelopmental disorder is readily modeled using SOSRS.

**Figure 8.**
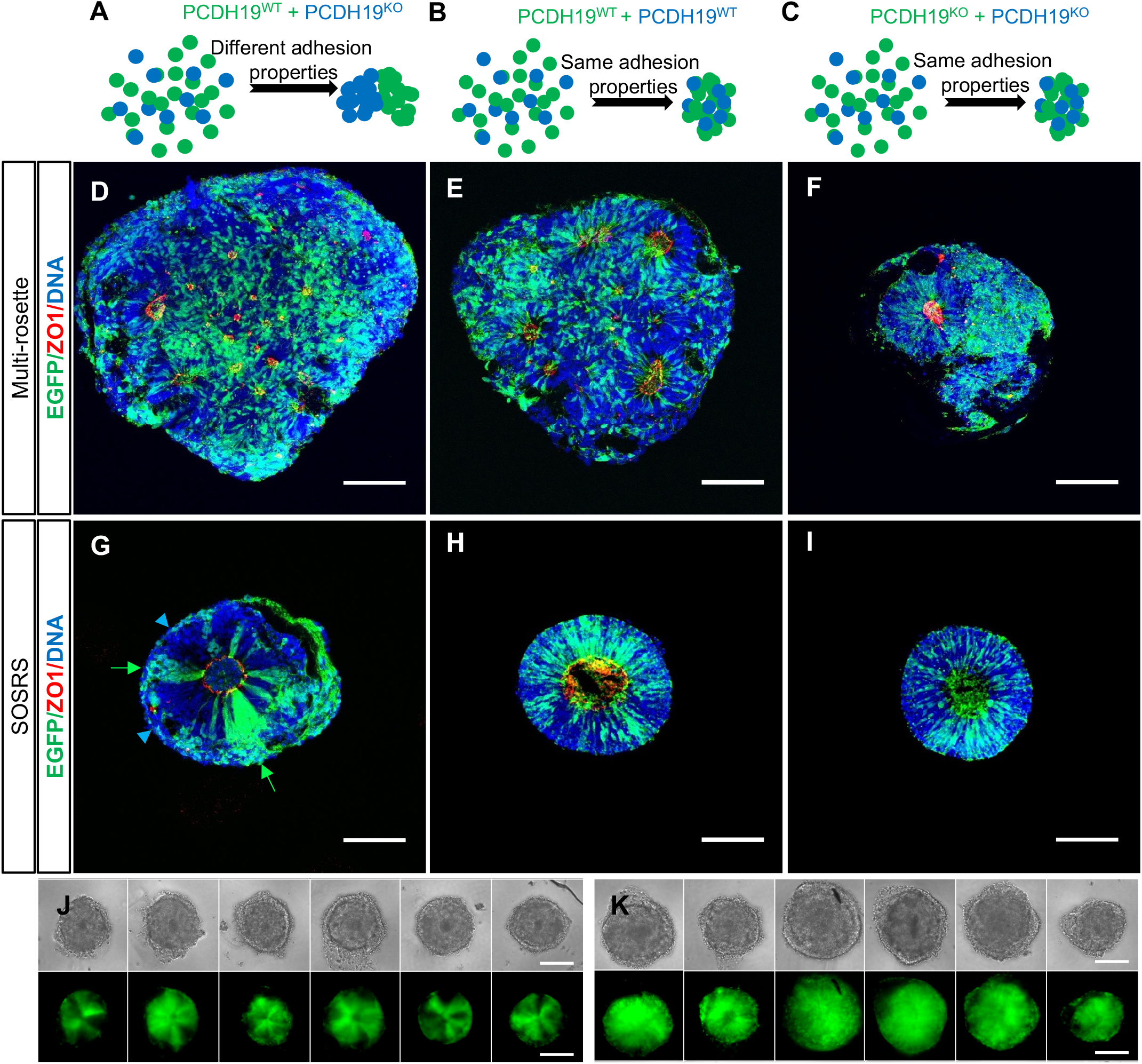
SOSRS recapitulate the early cell segregation phenotype found in mouse models of PCE. (**A-C**) Schematics depicting the hypothesized structural outcomes of brain organoids derived from different mixtures (simulating mosaicism of random X-inactivation) of GFP+ and unlabeled isogenic WT and KO cell lines based on previous data from mouse models (Hoshina et al., 2021; Pederick et al., 2018). (**D-F**) Confocal micrographs of multi-rosette brain organoids generated according to previously published methods (Dang et al., 2021; Qian et al., 2016) for each of the three genotype mixtures: WT/KO (**D**), WT/WT (**E**), and KO/KO (**F**). (**G-I**) SOSRS were generated in parallel with the same three mixtures: WT/KO (**G**), WT/WT (**H**), and KO/KO (**I**). Green arrows and blue arrowheads (in **G**) indicate SOSRS with segregated stripes of only WT or only KO cells. All organoids were immunostained for GFP (green) and ZO-1 (red), with bis-benzimide nuclear stain (blue). (**J**,**K**) Six individual SOSRS per condition from separate batches with GFP+ WT/GFP-KO mixtures (**J**) showing a clear segregation phenotype between the two isogenic cell populations, and GFP+ WT/GFP-WT mixtures (**K**) showing no segregation phenotype. All images are from day 20 spin-Ω organoids (**D**-**F**) or SOSRS (**G**-**K**). Scale bars are 100 µm.

## DISCUSSION

Here we describe simple, efficient production of a SOSRS brain organoid model with dorsal forebrain identity. We show that SOSRS have reproducible growth kinetics, VZ/SVZ-like structures, cortical lamination, and cellular diversity over five months in culture. SOSRS also demonstrate robust, reproducible structural abnormalities after chemical “teratogen” exposure or in modeling a genetic neurodevelopmental disorder.

A previous method to generate single rosette brain organoids involved geometric restriction in a micropatterned substrate (Knight et al., 2018). Our SOSRS model is produced without this complicated approach and can be grown for at least five months in culture with continued maturation and lamination of cortical-like structures. Several features of our protocol likely underlie the high reproducibility and fidelity of the SOSRS compared with previous brain organoid techniques. First, patterning likely occurs more efficiently in a cell monolayer than a 3D EB because of equal access of the entire monolayer to the patterning molecules and nutrients in the media. The reproducible size and growth of SOSRS probably reflects the size constraint imposed by monolayer cutting. Lamination and cell diversity may also be positively affected by the initial single rosette structure resulting in only one radial axis that can continue expanding in all directions rather than restriction from surrounding rosettes.

Unlike previous methods, SOSRS follow a normal developmental transition from a 2D organization to a 3D neural tube-like structure. The ability to monitor the exact timing of neurulation and lumen structure in whole-mount or live SOSRS allows for simple investigation of compounds that lead to NTDs. Exposure to the Rho-kinase or non-muscle myosin inhibitors (Y-27632 and blebbistatin) resulted in larger, more variably sized apical endfeet surface areas leading to dysmorphic lumens. In the context of *in vivo* neurodevelopment, these structural abnormalities would likely result in NTDs and, in fact, mouse *in utero* exposure to either compound results in severe NTDs (Escuin et al., 2015; Wei et al., 2001). Furthermore, an antiseizure medication that is teratogenic in humans, valproic acid, resulted in the same lumen enlargement while its non-teratogenic derivative, valnoctamide, did not. Thus, SOSRS should prove useful for studying mechanisms underlying NTDs and for CNS teratogen screening. Similarly, SOSRS revealed a robust morphological phenotype in modeling a genetic neurodevelopmental disorder. The early segregation phenotype in PCE has previously only been shown in a mouse model (Hoshina et al., 2021; Pederick et al., 2018). We were able to recapitulate this mosaic cell segregation pattern robustly in SOSRS but not using a multi-rosette cortical organoid method^7^. This finding suggests that the single rosette nature of SOSRS greatly increased our ability to adequately observe these early structural differences.

Reproducibility of the SOSRS structure and cell diversity has the potential to decrease the heterogeneity that impedes detailed pathogenic studies in the brain organoid field. We have shown that SOSRS have a remarkable ability to model early neurological disorders at the stages of neurulation and cortical mantle formation. Therefore, the use of this model system should provide improved fidelity in recapitulating human cortical development and reducing a major source of variability in disease modeling studies with existing brain organoid strategies.

## MATERIALS AND METHODS

### Human iPSC model

Control iPSC lines (Con2e, FF-H11, FF-E9, *PCDH19-*WT) were generated from commercially available foreskin fibroblasts as reported previously (Tidball et al., 2017) and were all male. A female control, CC-1, was obtained from Dr. Aaron Bowman (Tidball et al., 2015). Male iPSC lines with CRISPR based indels in the *PCDH19* gene (*PCDH19-*KO) were generated by simultaneous CRISPR-Cas9 gene editing and cellular reprogramming as reported previously (Tidball et al., 2017; Tidball et al., 2018). Another iPSC line used in this study, also a male line, is AICS-0023 in the Allen Cell Collection obtained from the Coriell Institute Biorepository. This line contains a monoallelic mEGFP-TJP1 (which encodes for the ZO-1 protein) to label the lumen in live SOSRS. Lastly, a clone of the H9 hESC line was produced with a 3.5 kb human DCX promoter driven mCherry using MiniTol2 vectors as previously described (Gupta et al., 2018). and is a gift from Dr. Michael Uhler. All of the experiments involving human cells were approved by University of Michigan IRB and Human Pluripotent Stem Cell Research Oversight (HPSCRO) Committee committees.

Cultures were maintained on Geltrex-coated 6-well tissue culture dishes in mTeSR1 medium at 37 °C prior to differentiation. When the colonies reached ∼40% confluency, the cultures were incubated with 1 mL of L7 dissociation solution for 2 minutes at 37 °C. The solution was then replaced with mTeSR1, scraped with a mini cell scraper, and pipetted up and down 3-6 times to break colonies into smaller pieces. The solution was then replated at a dilution of 1:12 onto newly Geltrex-coated dishes. Genomic integrity was accessed by high density SNP chip microarrays to identify CNVs. PCR for mycoplasma DNA was also regularly performed.

### SOSRS differentiation

iPSC lines were passaged using Accutase (Innovative cell) and replated onto Geltrex-coated (1:50 dilution in DMEM/F12) 12-well plates at 3-4 × 10^5^ cells/well in mTeSR1 with 10 µM rho-kinase inhibitor (Y-27632; Tocris, 1254). Medium without the inhibitor was replaced daily until the cells reach 80-100% confluency. The medium was then changed to 3N (50:50 DMEM/F12:neurobasal with N2 and B27 supplements) (Shi et al., 2012) without vitamin A with 2 µM DMH1 (Tocris, 4126), 2 µM XAV939 (Cayman Chemical, 13596), and 10 µM SB431542 (Cayman Chemical, 13031) (2 mL of media per well). Daily 75% media changes with 1 µM cyclopamine (Cayman Chemical, 11321) added began on day 1. On day 4, the monolayer was cut into squares using the StemPro EZ passage tool, and 1 mL of conditioned media was reserved. The squares were incubated for 1 minute with L7 hPSC passaging solution (Lonza). After aspirating the L7 solution, the squares were sprayed off the bottom of the culture plate with the 1 mL of preconditioned culture media with a P1000 micropipette. An additional 2 mL of fresh culture medium with the 4 inhibitors was added for a final volume of 3 mL. Approximately 200 µL of resuspended squares were then transferred into each well of a 96-well plate preincubated at 37 °C for 20 minutes with 35 µL of 100% Geltrex solution/well. Up to 16 wells can be made using one monolayer. Higher densities will result in fusion between nearby SOSRS forming multi-rosette doublets or “strings”. After 48 hours, daily 50% media changes were performed. On day 6 (2 days after cutting), the four inhibitors were no longer added but replaced with 3 µM CHIR99021. After 3-4 additional days of culture. The organoids were individually removed using the STRIPPER Micropipetter (Cooper Surgical) with 275 µM tips and plated individually in the wells of a low-adherence U-bottom 96-well plate with 200 µL of 3N medium without vitamin A with 3 µM CHIR99021, BDNF (20 ng/mL), and NT3 (20 ng/mL). The ideal diameter for picking the SOSRS is ∼ 250 µM. Starting 2 days later, half-media changes of 3N with vitamin A, BDNF (20 ng/mL), and NT3 (20 ng/mL) were repeated every other day. After 35 days of differentiation, 3N media with vitamin A (but without BDNF and NT3) was used for half-media changes every other day. At this time, SOSRS were transferred from the low-adherence 96-well plates to low-adherence 24-well plates for a necessary increase in media volume due to expanded SOSRS size.

### Immunocytochemistry

SOSRS imaged before day 14 were grown on Nunc™ Lab-Tek™ III 8-well chamber slides (Thermo Scientific) and fixed in paraformaldehyde for 30 min at room temperature (RT). Older SOSRS were fixed for 1 hour in 4% paraformaldehyde in PBS in suspension at 4 °C. After 3 PBS washes, the SOSRS were incubated in 30% sucrose overnight at 4 °C followed by embedding in TFM medium and freezing on dry ice. The blocks were sectioned on a Cryostat at 20 µM thickness. After 2 hours of drying at RT, the sections were washed 3 times with PBS to remove the TFM medium. For both formats, cells were permeabilized with 0.2% Triton-X 100 for 20 min at RT followed by incubation in PBS containing 5% normal goat serum with 1% BSA and 0.05% Triton-X100 for 1 h at RT. All samples were incubated in primary antibody overnight (see **Table S5** for antibodies and dilutions) in the same blocking buffer at 4 °C, washed 4 times in PBS with 0.05% Tween-20 (PBST), and incubated for 90 min with secondary antibody. Cells were washed 3 times in PBST and incubated with bisbenzimide for 5 min. After additional PBS washes, coverslips were mounted on slides with Glycergel mounting medium (Agilent Dako). Images were obtained on a Leica SP5 upright DMI 6000 confocal microscope.

### scRNA-seq data collection

One-, three-, or five-month SOSRS were dissociated using the Worthington papain dissociation kit. One mL of papain/DNAse solution was used for each organoid in a low-adherence 24-well plate. Organoids were incubated at 37 ºC for 45 minutes with gentle orbital shaking followed by 10 hard passes through a P1000 micropipettor. Organoids were incubated for an additional 15 minutes followed by 5 additional passes through a P1000 tip. Each organoid was labeled with sample-specific hashtag-labelled antibodies (HTO-A, Biolegend) to allow multi-sample pooled analyses to reduce batch effects, with subsequent de-multiplexing of scRNA-seq data based on the hashtags. Cell suspensions were passed through a 35 µM filter, and cells from multiple organoids were then mixed in equal quantities before submitting to the University of Michigan Advanced Genomics Core. Cell viability was found to be greater than 90% in all samples by Trypan blue exclusion. Single-cell suspensions were processed using the 10X Genomics Chromium platform, and scRNA-seq libraries were sequenced by Illumina NovaSeq. Each paired-end read contains a cell barcode and a UMI in Read 1, and the transcript-specific read in Read 2 with a read length of 151 nt. Separately, each cell generates an HTO count vector for the hashtags used, allowing the cells to be assigned to organoid samples they come from.

### scRNA-seq data analyses

#### Data processing and filtering

The transcript reads were aligned to human reference genome GRCh38, and the barcodes and UMIs were processed by *Cell Ranger v4*.*0*.*0*, which generated single-cell gene expression count matrices for each sample. We included cells that passed the cell size factor filter of >500 detected genes and the cell integrity filter of <10% mitochondrial transcripts. The HTO data were processed using *CITE-seq-Count v1*.*4*.*4* with default parameters. Cells present in both the filtered gene expression data and the HTO counts data were kept. For each cell, we normalized the raw transcript counts by (1) dividing by the total number of UMIs per cell and (2) multiplying by 10,000 to obtain a transcripts-per-10K measure, and then log-transformed by E=ln(transcripts-per-10K+1). For some downstream analyses, the normalized gene expression matrix was standardized for genes by centering and scaling for each gene using (E-mean(E))/sd(E).

#### Assignment of cells to samples based on HTO enrichment

We initially processed the gene expression and HTO counts data using the R package *Seurat v3*.*2*.*3*. Raw HTO counts were normalized for each cell using centered log ratio (CLR) transformation: for a given HTO, the counts in individual cells were divided by the geometric mean across cells and then log-transformed. We first applied the *HTODemux()* function in *Seurat*, but found the results unreliable, mainly due to the fixed cutoff of q=0.99 to designate the cells as positive or negative for an HTO. We modified the procedure by examining the joint distributions of the normalized HTO counts for multiple HTOs. For each HTO, we fit its distribution with two Gaussian curves, using the intersect point between the two modes as the optimal cutoff. Cells that were positive for two or more HTOs were classified as doublets and removed from further analyses. Cells negative for all HTOs were also removed. Compared to the default method, our modifications recovered more singlets (**Fig. S8A-C**). In all, we identified 4,027 singlets with an average of 4,214 detected genes and 15,216 UMIs for the 1-month old data, and 9,314 singlets with an average of 4,528 detected genes and 16,595 UMIs for the 3-month old data.

#### Clustering and visualization

Since the 3-month old data contained 2 samples, each with 3 HTO tags, we corrected for potential batch effects using *IntegrateData()* function in Seurat. For visualization of HTO data, we used Euclidean distances calculated from the normalized HTO data as inputs for tSNE (**Fig. S8D-G**), and colored cells based on their HTO-based assignment described above. For visualization and clustering of gene expression data, we performed principal component analysis (PCA) on the standardized gene expression matrix, selected top PCs based on elbow position of scree plot, and did tSNE, UMAP and Louvain-Jaccard clustering using top PCs. To evaluate the similarity of identified clusters among samples, we examined (1) the proportional constitution to different clusters from each HTO tag (**Fig. S4B**,**C**), and calculated the rank correlation of cluster centroids for the clusters found in different HTO-tagged samples. Cluster-specific markers were obtained by comparing cells in a given cluster with those in all other clusters using the binomial likelihood test. Selection criteria were (1) at least 20% difference in detection rate; (2) a minimum of 2-fold higher mean expression level in the cluster compared to all other clusters.

#### Annotation of cell types

We annotated the clusters based on sets of highly informative markers for known brain cell populations, cluster-specific markers produced from our analysis, and comparisons with other brain organoid reference datasets: the brain and organoid atlas (Tanaka et al., 2020), and the cortical organoid atlas (Bhaduri et al., 2020).

### Electrophysiological recordings

Electrophysiological recordings were performed on neurons from 3-month SOSRS that were dissociated and cultured adherently for 35 days on PEI-laminin-coated MatTek dishes. Excitatory neurons were labeled with lentiviral CAMKIIα-driven GFP (Tidball et al., 2020). Recordings were made at RT in BrainPhys medium. Individual coverslips with plated cells were transferred into the recording chamber and continuously perfused with BrainPhys bubbled with a mixture of CO_2_ (5%) and O2 (95%). Neurons were identified for patch clamp recordings using a 40x water-immersion objective and infrared differential interference contrast (IR-DIC) optics that were visualized using NIS Elements image analysis software (Nikon). Data were acquired using an Axon CV-7B headstage amplifier, Axon Multiclamp 700B amplifier, Digidata 1440A interface, and pClamp-10 software (MDS Analytical Technologies). All voltage data were acquired in the current-clamp mode with bridge balance compensation and filtered at 2 kHz.

Borosilicate glass patch pipettes were pulled with resistances of 3-5 MΩ. Patch electrodes were filled with internal solution containing the following (in mm): 119 K Gluconate, 2 Na gluconate, 6 NaCl, 2 MgCl2, 0.9 EGTA, 10 HEPES, 14 Tris-phosphocreatine, 4 MgATP, 0.3 tris-GTP, pH 7.3, osmolarity 290. Voltages were not corrected for liquid junction potential (+10 mV). Electrode capacitance was compensated on-line in the cell-attached mode. To control the quality and stability of recording, access resistance was monitored continuously during recording, and data were discarded if the access resistance rose above 40 MΩ or changed by >20%. Exemplar traces were plotted using Prism (GraphPad).

### SOSRS drug treatment

SOSRS were generated as described above on either 96-well plates or LabTek chamber slides. For the Y-27632 and blebbistatin experiments, one day following addition of monolayer fragments to Geltrex, 50% of the media was exchanged (100 µL) for fresh media containing either vehicle (DMSO), rho-kinase inhibitor (Y-27632), blebbistatin, valproic acid, or valnoctamide. After an additional 48 hours of incubation, the SOSRS were fixed and immunostained as described above. For the valproic acid experiments, the ZO-1-EGFP expressing monolayer fragments were treated immediately after plating for 48 hours followed by fixation and staining with bis-benzamide only.

### *PCHD19* iPSC validation and organoid culture

A male *PCDH19* KO iPSC line containing a 2 bp deletion and isogenic control were generated previously (Tidball et al., 2017). To confirm that the 2 bp deletion predicted to cause a loss-of-function frame shift mutation resulted in nonsense mediated decay, we performed quantitative reverse transcriptase PCR (qRT-PCR) to measure PCDH19 mRNA expression (**Fig. S7A**,**B**). Total RNA was isolated with the miRNeasy Mini Kit (Qiagen) according to the manufacturer’s instructions. RNA (500 µg) was used to generated cDNA using the SuperScript III First-Strand Synthesis SuperMix kit (Invitrogen #11752-050). qRT-PCR was carried out with Power SYBR Green PCR Master Mix (Applied Biosystems, #4367659). Two pairs of primers were used to amplify two different exonic regions of the PCDH19 transcript (5’ end, and 3’ end) respectively. Fold changes were calculated using 2^-ΔΔCt^ method. Primers can be found in **Table S6**. We knocked in a ubiquitous GFP reporter into the AAVS1 (safe harbor) locus in each of *PCDH19* isogenic WT and KO lines using transcription activator-like effector nucleases (TALEN). The expression of GFP was confirmed by both fluorescence-activated cell sorting and microscopy. Plasmids used were: AAVS1-CAG-GFP, AAVS1-TALEN-L, AAVS1-TALEN-R. AAVS1-CAG-hrGFP was a gift from Su-Chun Zhang (Addgene plasmid # 52344) (Qian et al., 2014). AAVS1-TALEN-L and AAVS1-TALEN-R were gifts from Danwei Huangfu (Addgene plasmid # 59025 and 59026) (González et al., 2014). Unlike all other lines used here, the *PCDH19* and isogenic control iPSC lines were grown in TeSR-E8 medium and passaged every 4-5 days by EDTA (0.8mM) and 10 µM ROCK inhibitor (Y27632) treatment.

To replicate the mosaic expression of *PCDH19* found in PCE patients (heterozygous females or mosaic males), we mixed GFP-labeled WT iPSCs with unlabeled isogenic KO *PCDH19* iPSCs at a 1:1 ratio as a “PCE mosaic” condition on the initial day of brain organoid differentiation. Similarly, we mixed cultures of GFP-labeled/unlabeled (1:1) KO only or isogenic WT only cells as controls. We simultaneously differentiated the three types of mixed iPSC cultures into cortical organoids using two different methods, each with the same starting number of cells (∼2×10^6^ per mix). The first method is the SOSRS protocol described above, with a minor modification of 1 µM CHIR treatment between days 6-10. The second method followed a published protocol known as Spin-Ω (Qian et al., 2016) which we have used previously (Dang et al., 2021). In brief, we generated EBs (day 0) from mixed iPSC cultures in AggreWell800 plates (STEMCELL Technologies). On day 1, the EBs were transferred to a low attachment 60 mm petri dish containing Dorsomorphin/A83 medium and grown for 4 days. On day 5, the EBs were gradually switched to CS induction medium (with SB-431542, CHIR, and Heparin). On day 7, about 20 EBs were embedded in Matrigel and then grown in one well of a 6-well plate for 7 days in CS induction medium. On day 14, the embedded EBs were transferred to a spinning mini-bioreactor (Spin-Ω) and cultured in B27 differentiation medium without Vitamin A. On day 20, organoids were fixed, embedded with O.C.T in cryomold, and stored at −80°C for eventual sectioning and immunostaining. Lastly, similar to the mixing experiment that was described for male iPSC lines, we generated red and green mixed SOSRS from a female PCDH19 KO hESC line and an isogenic WT line, both of which stably expressed GFP or RFP using the lentivirus integration approach.

### Lumen image analysis and quantification

ImageJ was used to subtract background from 63x confocal images and create a binary mask over the ZO-1 image. The “fill hole” function was used to create a filled mask, and both area and circularity were measured using bis-benzamide nuclear stain to identify total SOSRS area and ZO-1-EGFP or immunolabelled ZO-1 to detect the apical lumen.

### Statistical analyses

Prism Graphpad was used for all statistical analyses. In **Fig. 1H**, a linear regression was performed to obtain an R^2^ value and slope. In **Fig. 5**, non-parametric analyses were used for all SOSRS lumen measurements based on Prism normality tests of the data sets. In **Fig. 5 D,E,H,L**, data was analyzed by Welch’s t-test for significance. Analysis in **Fig. 5 L,M** was performed using Kruskal-Wallis due to multiple group comparisons. All other statistical details can be found in the figure legends.

## ACKNOWLEDGEMENTS

Part of this research was made possible through the Allen Cell Collection, available from Coriell Institute for Medical Research. We would like to thank Michael Uhler, Ranmal Samarasinghe, and M. Carmen Varela who provided important manuscript feedback.

## Data and code availability

- All single-cell mRNA sequencing data of this study have been deposited in the Gene Expression Omnibus (GEO) under reviewer access at: https://www.ncbi.nlm.nih.gov/geo/query/acc.cgi?acc=GSE181518 Reviewer token: kpmnqsyalzchzmz
- All original code has been deposited at Github and is publicly available as of the date of publication. https://github.com/qianqianshao/SOSRS
- Any additional information required to reanalyze the data reported in this paper is available from the lead contact upon request.

